# Detecting movement preparation with HYDRE: a hybrid model of temporal and spectral brain signals

**DOI:** 10.64898/2026.10.14.758540

**Authors:** Romaric Thiboud, Fayed Rassoulou, Denis Schwartz, Pauline Mouches, Pierre-Elliott Thiboud, Eshaan Eshaan, Alessandro Farnè, Marine Vernet

**Affiliations:** IMPACT Team of the Lyon Neuroscience Research Center (CRNL) CNRS (UMR 5292) INSERM (U1028) University Claude Bernard Lyon (UCBL), Lyon, France; COPHY Team of the Lyon Neuroscience Research Center (CRNL) CNRS (UMR 5292) INSERM (U1028) University Claude Bernard Lyon (UCBL), Lyon, France; CERMEP-Imagerie du Vivant, Lyon, France; EDUWELL Team of the Lyon Neuroscience Research Center (CRNL) CNRS (UMR 5292) INSERM (U1028) University Claude Bernard Lyon (UCBL), Lyon, France; MYRIAD Team Centre de Recherche en Acquisition et Traitement de l’Image pour la Santé (CREATIS), INSA Lyon, University Claude Bernard Lyon (UCBL), CNRS (UMR 5220), INSERM (U1294), Lyon, France; Neuro-Immersion Platform of the Lyon Neuroscience Research Center (CRNL) CNRS (UMR 5292) INSERM (U1028) University Claude Bernard Lyon (UCBL), Lyon, France

## Abstract

Previous research on brain-computer interfaces (BCIs) has demonstrated the feasibility of decoding different movements. However, less attention has been given to detecting motor preparation (i.e., decoding from brain signals whether or not a participant is about to perform a given movement). In this study, we wanted to assess the performance of a new model, rooted in motor preparation physiology, in detecting an upcoming movement across the period preceding a voluntary action.

Participants (n=18, including 7 females) performed a simple self-paced motor task where they either pressed a button to change an image shown on screen, or let the computer change the image. Participants’ brain activity was recorded using magnetoencephalography and submitted to a newly developed linear ensemble classifier that relies on features extracted using PCA from the temporal, spectral and Riemannian domain. We then compared its performance against other classifiers used in BCIs.

Our HYbrid Domain Riemannian Ensemble (HYDRE) classifier emerged as the best-performing method, achieving average above-chance decoding up to 1400ms before the action, with a maximum ROC score of 76% at the time of the button press. Moreover, decoding performance in the temporal and Riemannian domains was respectively predictive of Readiness Field and Event-Related Desynchronization magnitudes, two well-known motor preparation markers.

Combining features from the temporal, spectral, and Riemannian domains significantly improved motor preparation decoding compared to single-domain approaches. By leveraging the most discriminative features from each domain, HYDRE yielded consistent performance gains across participants with varying baseline decoding performance, effectively addressing inter-individual variability in motor preparation markers.

**Significance Statement:** Our results show that combining features from temporal, spectral, and Riemannian domains significantly improves motor preparation decoding compared to single-domain approaches. The HYbrid Domain Riemannian Ensemble (HYDRE) classifier we developed following this principle yields consistent performance gains across participants with varying levels of baseline decoding performances (improving prediction in both high- and low-decoding individuals). Lastly, we characterize how motor preparation decoding evolves in the time window leading up to the movement, linking decoding trajectories directly to observable physiological markers.

## Introduction

Brain-computer interfaces (BCIs) are systems that establish real-time communication between neural activity and external devices (through computer) by decoding brain signals and translating them into control commands (Ramadan & Vasilakos, 2017). Many motor-related BCI studies rely on motor imagery paradigms and focus on binary classification, such as distinguishing between imagined left-versus right-hand movements (Huang et al., 2009), or extending to multiple effectors (Wang et al., 2012). This kind of BCI often performed very well due to the asymmetric nature of motor markers in electrophysiological signals (Shibasaki & Hallett, 2006). More challenging, predicting upcoming movements from pre-movement brain activity could help to predict a user’s intended outcomes (Beste et al., 2026), and improve already working motor imagery based BCI (Y. Zhang et al., 2025).

Previous attempts to detect motor preparation from EEG time series have reported promising results. When classifying trials with vs. without motor preparation, Ody et al. (2023) achieved above-chance decoding as early as 800 ms before a button press, peaking at 85% average accuracy at press. Bai et al. (2011) detected movement preparation in 40% of self-paced wrist extensions (75% precision), on average 620 ms before execution (EMG onset). More recently, in a series of 3 articles Schultze-Kraft et al. (2016, 2020, 2021) applied regularized linear discriminant analysis (or common spatial patterns) to epoched signals from electrodes with strong preparatory activity, detecting upcoming EMG onset in up to 55.9% of trials (56.6% precision) in their early BCI and reaching 80% offline accuracy in their later study (Schultze-Kraft et al., 2020). Overall, motor preparation can be decoded, albeit with high inter-individual variability, but few studies describe how decoding performance evolves in the period preceding movement onset.

Several electrophysiological motor preparation markers have been described in the literature. In the temporal domain, the Readiness Potential (RP) in EEG (Kornhuber & Deecke, 1965) or Readiness Field (RF) in MEG is a slow negative deflection over motor areas that begins over 1 s before self-initiated action and peaks at movement onset. It has long been established that the RP exhibits large inter- and intra-individual variability in both shape and amplitude (Libet et al., 1982, 1983). Besides RP, another maker of motor preparation is the Event-Related Desynchronization (ERD) visible in the spectral domain. ERD is characterized by a decrease of alpha and beta band power (8-30 Hz) during movement preparation and execution (Shibasaki & Hallett, 2006). Studies tends to agree that RP and ERD represent different neural processes, even if the exact function of each of them is still debated (Parés-Pujolràs et al., 2023; Shibasaki & Hallett, 2006; Wairagkar et al., 2018). If these two processes encode complementary information about motor preparation, combining features from both the temporal and time-frequency domains may enhance the detectability of motor preparation. This approach of combining features from different domains has been already applied to improve the left vs. right hand motor imagery decoding (Corsi et al., 2022; Radman et al., 2021) and even, to some extent, for motor preparation (Ieracitano et al., 2021).

Here, we propose a linear classifier, the HYbrid Domain Riemannian Ensemble (HYDRE) classifier to improve motor preparation decoding. This model relies on temporal and spectral domain features to predict the motor preparation of self-paced button presses. In this study, we evaluate the HYDRE classifier using offline, single-trial classification, comparing its performance against state-of-the-art models across the motor preparation phase. We place particular emphasis on a window of interest from −300 ms to −250 ms prior to the button press. This window is close to the ‘point of no return’ beyond which a planned movement can no longer be canceled (Schultze-Kraft et al., 2016), while remaining early enough to compensate for inherent BCI processing latencies. Enhancing decoding performance during this period could improve BCI-assisted interventions, for example, by facilitating stroke recovery through the initiation of a movement aligned with the patient’s motor intent to execute it.

## Methods

### Participants

18 healthy participants aged between 19 and 33 years (7 women; mean = 24 +/− 4 years) were recruited for this experiment. All participants reported no history of neurological or psychiatric conditions, had normal or corrected-to-normal vision, provided written informed consent before taking part in the study, and received a monetary compensation of 60 euros for their participation. The experiment was validated by a Institutional Review Board (CPP sud-est IV 2010-A01180-39) in accordance with the Declaration of Helsinki. The experiment was designed in 2021 and the data collected between March and November 2021.

### Material

Participants were seated in a MEG system (CTF Omega 275) inside a magnetically shielded room. The MEG signal was acquired using 274 axial gradiometers, a right temporal sensor (MRT36) being broken. An eye tracker (Eyelink 1000, SR Research) was used to track gaze position and pupil size, and monitor eye blinks throughout the experiment. The task, programmed in Python (version 3.6) with Psychopy (Peirce et al., 2019), controlled the visual stimuli presentation on a screen. Triggers were sent via a parallel port to the MEG recording computer, which also synchronised button presses, as well as the signals from a photodiode, ensuring precise timing with the visual stimulation.

Visual stimuli were displayed using a PROPixx projector (VPixx Technologies) and a 40 by 23 cm screen, with a 1920 × 1080 pixel resolution and a 120 Hz refresh rate. Images adapted from the COCO dataset (Lin et al., 2014) were used as visual stimuli. An initial pool of neutral images depicting ordinary situations was collected to ensure comparable salience and content across the set. Images were processed in MATLAB according to the following criteria: colored images with a minimum size of 480 × 480 pixels were retained and resized to 480 × 480 pixels if needed. To prevent cropping the main element, an algorithm recentered each image on its center of mass prior to resizing. Brightness was normalized to the mean brightness of the full set. Finally, all images were visually inspected, and those containing a central face, stop signs, vulgar content, whose normalization rendered them too dark, too bright, or ill-cropped were excluded. From this pool, 1400 images were randomly selected (allowing up to 7 blocks of 200 images).

### Experimental design

Participants performed a self-paced action task in which naturalistic images were presented one after the other at the center of a screen located 64 centimeters away from them (***Fig. 1***). At the center of each image, a red fixation cross (0.8 cm in width and height) was presented and participants were asked to fixate the cross during the entirety of the block, while avoiding blinking as much as possible. Following the presentation of an image, participants were instructed to press a button in order to proceed to the next image whenever they felt they had observed the image long enough to be able to describe it. Participants were also informed that the computer could change the image at a random time. To have similar trial duration between participant generated and computer generated trials, this time was sampled from a gamma distribution (k=3, θ=1) that closely matched the button press time distribution of pilot participants. The gamma distribution was also cut to only generate value superior to 500 ms. Each button press (later defining “human” trial) or automatic switching to the next image (later defining “computer” trial) was followed 20-30 ms later by a gray screen lasting on average 24ms ± 1.2ms before the new image appeared on screen. Finally, on each break, participants received feedback encouraging them to press the button more often or less often, depending on the respective number of “human” and “computer” trials, ensuring a similar number of trials for each category.

**Figure 1:**
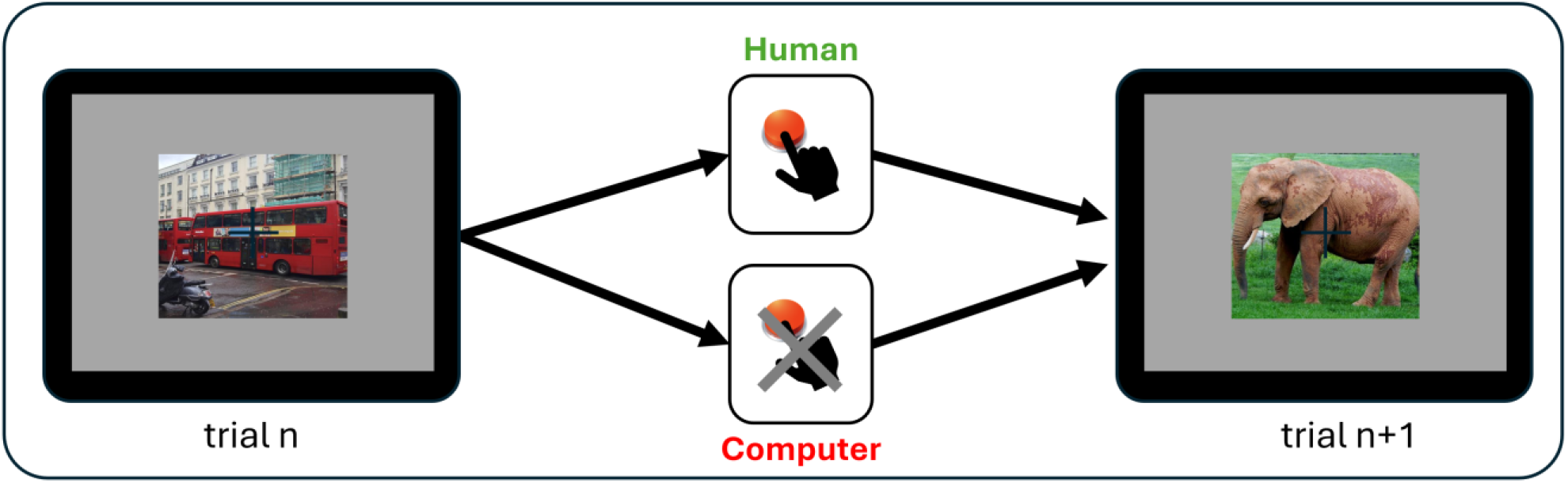
Self-paced action task. Participants were presented with images and were instructed to press a button whenever they felt that they observed the image long enough. The computer could also change the image. At the end of the block participants were encouraged to press the button “more often” or “less often” depending on the number of trials in which the image change was due to themselves or to the computer, with the aim of balancing the two types of trials.

During the session at least 1200 images (600 “human” trials and 600 “computer” trials) were presented to the participants divided in blocks of 200 trials spaced by a resting period during which the participant’s head position in the MEG was checked. Blocks were further divided in 50 images sub-blocks separated by 3 short breaks. If the participant moved too much (>5mm) during a block, the corrupted block was discarded from further analysis and replaced by additional subblocks of 50 trials until at least 600 “human” trials and 600 “computer” trials were completed.

Participants performed the task using their right hand and a button-box with 2 buttons. They ended the short breaks inside each block by pressing the right button with their middle finger, while they processed to the next trial by pressing the left button with the index finger.

### MEG pre-processing

MEG signals were initially acquired at a sampling rate of 600 Hz and subsequently downsampled to 120 Hz. For the last 8 participants an online forward band-pass filter (0.1–30 Hz) was directly applied to the data. On the continuous data, if two commands triggering an image change were spaced by less than 500ms the second one was discarded to ensure enough time for the motor preparation decoding. A forward–backward FIR band-pass filter (0.1–30 Hz) was applied to the continuous data, and epochs were extracted relative to the command triggering an image change, i.e., the button press (either the human button press or the “virtual” button press, i.e., the computer command), using a −2s to +1s window. Because the multivariate analyses focused on motor preparation and to reduce computational load, decoding was performed on a restricted time window ranging from −1.5s to 0s. To stay as close as possible to the BCI framework no further pre-processing was applied on data used to train the different classifiers.

For univariate analysis, correction of cardiac and ocular artifacts based on typical scalp topography and time course was performed on epochs using an independent component analysis (ICA) (Jung et al., 2000). After manual artefacts rejection with the ICA (mean number of ICA rejected by subject: 3.67±1.34 out of 100 independent components estimated with “fastICA” (Hyvarinen, 1999), epochs were submitted to an algorithm from the autoreject toolbox (version 0.4.3) (Jas et al., 2016, 2017) that automatically interpolates bad segments (i.e defined as a single sensor within an epoch), or reject bad epochs (mean percentage of interpolated segments: 0.87±0.36%; mean percentage of rejected epochs: 1.69±2.09%)

### Analysis

All the data analyses presented below were carried out using Python 3.12.

#### Classification models

In this study, we applied classifiers at each time point and, unless otherwise specified, classification methods were implemented using default parameters as provided in scikit-learn (version 1.7.1). Models were trained on normalized brain signals (z-score standardization) using the standard scaler function from the scikit-learn toolbox (version 1.7.1). Normalisation was applied per participant, per fold, with the mean and standard deviation of the brain signal computed across trials per sensor and time window. Evaluation was performed through a stratified 10-fold cross-validation procedure. In each fold, standardization and feature extraction parameters (e.g., Principal Component Analysis (PCA), Tangent space projection for the Riemannian module, CSP, etc.) were computed on training data only and later applied to the entire dataset. All models were trained and applied over the last 1500 ms leading to the button press or the computer command to change the image, resulting in 30 time points per model tested. All models, except the CSP+LDA classifier, Riemannian and Spectral modules, were applied on the last 50-ms preceding each time point. The time window for the CSP+LDA and Riemannian module was increased up to the last 200-ms in order to improve the stability of the covariance matrix estimation. The time window for the Spectral module was increased up to the last 500 ms in order to accurately compute power in the alpha band, before power being extracted from the last 200 ms.

##### HYbride Domain Riemannian Ensemble (HYDRE) classifier

The HYDRE model is grounded in the ensemble learning theory, which asserts that the aggregation of multiple models produces a better collective decision to that of any single constituent model. In our case the classifier extracts features from the temporal, Riemannian and spectral domain to perform classification, using logistic regression, in each domain separately. The HYDRE classifier is composed of 3 independent modules that are later ensembled (***Fig. 2***). Classification performance of each module is evaluated in addition to those of the complete model.

**Figure 2:**
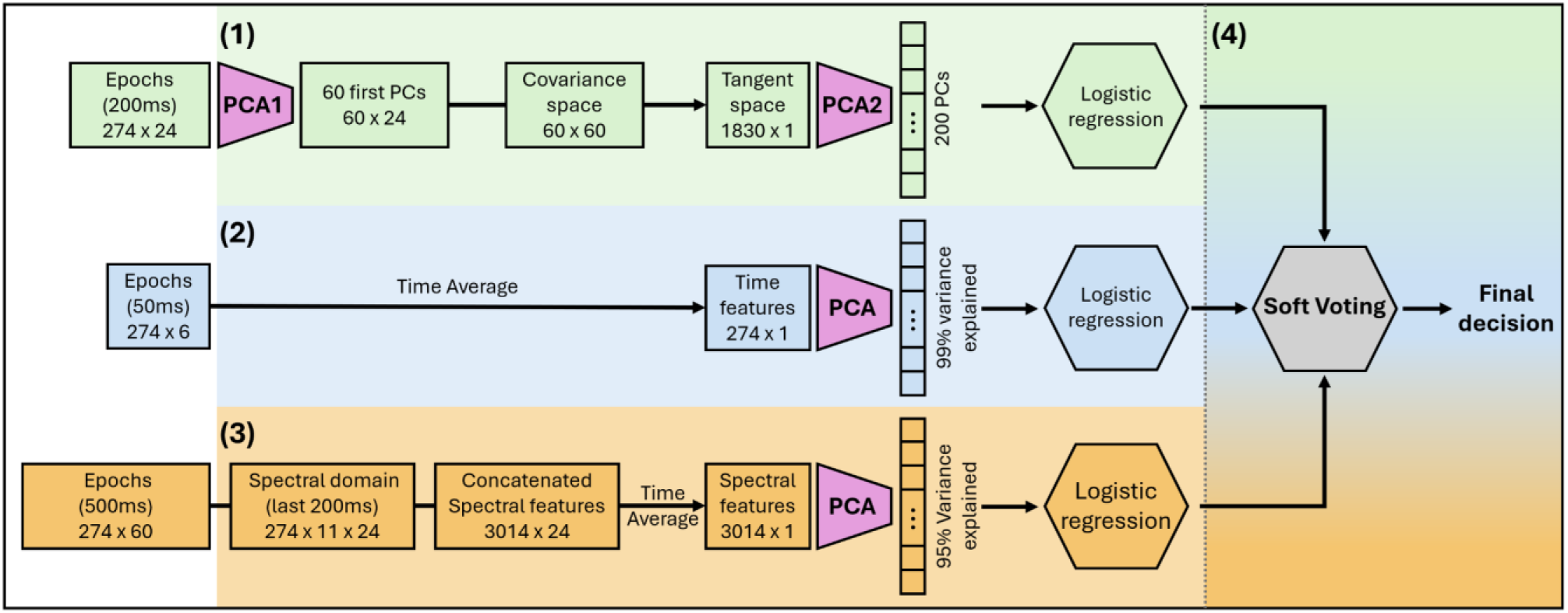
The process of HYbride Domain Riemannian Ensemble (HYDRE) classifier. **(1)** Riemannian module (light green): 200ms of band-pass filtered (0.5-30 Hz) brain signal encompassing the 50ms window of interest was used (see methods for details). A first PCA was applied to reduce dimensionality by selecting the first 60 principal components. The covariance matrix was then computed using OAS regularization and the data was then projected on the tangent space using the log-euclid rule. A second PCA was applied to select the first 200 principal components that were used as features to train a logistic regression. **(2)** Temporal module (light blue): 50ms of band-pass filtered (0.5-30 Hz) brain signal was used as input. Data were then averaged on the time dimension and dimensionality was reduced by applying a PCA and selecting principal components explaining 99% of the original variance as features for a logistic regression. **(3)** Spectral module (light orange): A time-frequency decomposition was performed using Multitaper on a 500ms window and then selecting the last 200ms of the time-frequency representation, encompassing the 50ms window of interest (see methods for details). Data were then concatenated over sensors and frequencies dimensions and averaged on the time dimension. A PCA was applied and principal components explaining 95% of the original variance were used as features for a logistic regression. **(4)** In a final step, the probability of the predicted label of the 3 modules were ensemble using soft voting to output the HYDRE classifier decision.

###### a) Temporal domain module

In this module, PCA was applied to the 274 sensors as a feature extraction step. The first *nt*ℎ principal components (PCs) explaining at least 99% of the total variance (representing an average of 53.57 ± 20.86 PCs across the time window of interest) were selected as features to train the model (***Fig. 2.2***). On each trial, the predicted probability of the “motor preparation” label was computed for use in the ensemble module.

###### b) Riemannian module

The principle of Riemannian geometry applied in BCI is that covariance matrices lie on a Riemannian manifold, not in ordinary Euclidean space. Working in this space provides meaningful properties, such as invariance to linear transformations and more robust averaging. But since most machine-learning methods assume Euclidean geometry, data need to be projected back to Euclidean space into a so-called tangent space, which is a local flat approximation of the manifold. The tangent space is typically defined at the Riemannian mean of the training covariance matrices. This point minimizes the sum of Riemannian distances to all samples, providing a statistically meaningful center of the data. This projection locally approximates the relevant geometric structure of the Riemannian manifold, while allowing the use of most classification methods.

Here, a first PCA was applied to extract the first 60 principal components to reduce dimensionality and optimize the speed of the covariance matrix estimation that was computed over a 200ms time window. To satisfy the Riemannian manifold requirements of symmetry and positive definiteness, Oracle Approximating Shrinkage (OAS) regularization was used during estimation. The resulting matrix was projected onto the tangent space using a log-Euclidean mapping. We then applied a second PCA, retaining the first 200 principal components, to extract relevant features of the tangent space. These components served as features for a logistic regression classifier with default hyperparameters (***Fig. 2.1***). All Riemannian geometry operations were implemented using the Pyriemann toolbox (v0.9).

###### c) Spectral domain module

In the last module, a Logistic Regression was trained at each time point of the time-frequency representation, which was estimated on a 500ms time window. To mitigate edge artifacts, a mirror padding of 250ms was added on both ends of the window resulting in a 1000 ms time window on which a Tukey window (α=0.83 ; N=60) was applied to progressively decrease the signal intensity out of the original 500ms time window. The time frequency representation was computed using a multi-taper spectral estimation (time-bandwidth=2) to extract power from the 8-30 Hz frequency band in 2 Hz increments, resulting in 12 frequencies. Then, padding was removed and the last 200ms of the original 500ms was selected. The sensor and frequency dimensions of the time-frequency representation were then concatenated and the data was averaged on the time dimension. These two steps reduced the data matrix from 4 dimensions (trials x sensors x frequencies x times) to 2 dimensions (trials x sensors*frequencies). Finally, a PCA was used as a feature extraction step: principal components explaining at least 95 percent of the total variance (representing an average of 140.74±72.96 PCs on the time window of interest) were selected as features to train a logistic regression (***Fig. 2.3***).

###### d) Ensemble the modules

For each trial, the predicted label probabilities (ranging from 0 to 1) from each module were aggregated into an ensemble following a soft voting rule (***Fig. 2.4***). If the averaged probability of the predicted label over the 3 modules exceeded a threshold of 0.5, “motor preparation” label was selected as the predicted class of the ensemble.

##### Standard classifiers

To comparatively evaluate the decoding performance of the HYDRE classifier and its constituent modules, three baseline classification methods were selected. First, standard Logistic Regression was included to quantify the performance gains achieved by manifold-based transformations and ensembling over a basic linear decision boundary (Lotte et al., 2018). Second, a Gradient Boosting classifier served as a non-linear ensemble benchmark to evaluate standard machine learning algorithms against the HYDRE classifier. Finally, Common Spatial Patterns combined with Linear Discriminant Analysis (CSP+LDA) was chosen as the traditional gold standard for motor BCI spatial filtering (Lotte et al., 2018).

###### a) Logistic regression classifier

A logistic regression classifier was trained in the temporal domain on normalized brain signals from all 274 sensors.

###### b) Gradient boosting tree applied on PCA extracted features

Gradient Boosting is an ensemble learning method that sequentially trains decision trees, where each new tree aims to correct the errors of the previous ensemble. This method allows modeling of complex, nonlinear relationships. Among methods using decision trees, we selected Gradient Boosting over random forest classifiers, because the former one builds trees sequentially rather than independently, often achieving higher accuracy albeit to a greater computational cost. A gradient boosting classifier (default parameters) was trained on temporal domain features extracted with a PCA (explaining more than 99 percent of the total variance).

###### c) Linear discriminant analysis (LDA) applied on Common Spatial Pattern (CSP) extracted features

Common Spatial Patterns (CSP) is a widely adopted feature extraction technique in BCI research, particularly for motor-imagery-based classification. CSP computes spatial filters that maximize the variance of one class while minimizing the variance of the other, resulting in discriminative log-variance features. In the present study, 20 spatial filters were extracted from the original data. The signals were not averaged over the 200ms time windows, since CSP relies on the covariance structure of the time samples to estimate the spatial filters. To improve the covariance estimation given the limited length of the time windows, OAS regularization was applied, resulting in increased numerical stability and reduced estimation noise. Then an LDA was applied on the extracted filters.

#### HYDRE classifier evaluation

To evaluate how the modules predictions are integrated to form the HYDRE classifier, Cohen’s Kappa was computed at each time window for every pair of modules (Riemannian/Temporal; Riemannian/Spectral; Temporal/Spectral) and averaged over the pairs. Cohen’s Kappa is bound between −1 and 1 and tells us how much the modules agree with each other beyond what would be expected by chance. A Cohen’s Kappa close to 1 indicates perfect agreement, the modules have identical predications. A Kappa value close to 0 means that the agreement is no better than if modules were guessing based on class frequencies. In the case of an ensemble model, a low Cohen’s Kappa indicates low redundancy between the modules.

#### Multivariate analysis

To evaluate models’ performance, the ROC AUC was computed for each 50 ms time window. To identify above-chance decoding of motor preparation signals, we performed non-parametric one sample cluster permutation t-tests independently for each method, using the default parameters in MNE-Python to correct for multiple comparisons. Furthermore, to identify significant differences in decoding performance between HYDRE and all the other classifiers, additional one-sample cluster-based permutation tests (MNE-Python toolbox with default parameters) were conducted for each classifier.

Temporal trends in decoding performance and modules’ average Cohen’s Kappa were assessed by first computing Spearman correlations at the participant level for each method or pair of methods. A group-level, one-tailed Wilcoxon signed-rank test (alternative: greater) was then used to determine if the resulting rho coefficients were significantly positive.

A focused analysis was conducted on the time window of interest (−300 ms to −250 ms before the button press). Group-level differences in variance between HYDRE and alternative classifiers were investigated using permutation tests (SciPy). Decoding performance differences between HYDRE and its modules were assessed using a Friedman test followed by Nemenyi post-hoc comparisons. Spearman correlations, corrected for multiple testing via the Holm-Bonferroni method, were used to evaluate group-level associations between the modules’ decoding performances, as well as between HYDRE and the best-performing module for each participant. Lastly, a one-sample t-test (alternative: greater) was employed to determine if the modules’ average Cohen’s Kappa significantly exceeded zero during this window.

To visually assess how well the HYDRE classifier and the other classifiers behave under different decision thresholds, the ROC curves were plotted for each method on the time window of interest. In addition, Youden’s J statistic was used to identify the optimal thresholds maximizing the difference between the true positive rate and the false positive rate.

#### MEG Univariate analysis

Electrophysiological data were compared between “computer” and “human” trials using univariate analyses across temporal and spectral domains to assess the presence of motor preparation markers, specifically examining Event-Related Fields (ERFs) and spectral power, respectively.

##### Temporal analysis

To investigate the Readiness Field, processed epochs (−2s before to 1s after the button press) were averaged separately for each condition (human and computer) at the participant level, and then used to compute the difference between conditions. Statistical significance at the group level was assessed using a one-sample spatio-temporal cluster-based permutation test performed on the between-conditions difference across all participants. The test was implemented using the default parameters in MNE-Python on a time window ranging from −2s to 0s before the button press.

##### Spectral analysis

For spectral analyses the tested window included the post-button press period (−2s before to 1s after the button press) to capture the ERD and the power resynchronisation following an action, the Event-Related Resynchronisation (ERS). The Time-Frequency Representation (TFR) was computed using a Morlet wavelet transform as implemented in the MNE-Python toolbox. For each epoch, spectral power was extracted for frequencies ranging from 8Hz to 30Hz in 1Hz increments. To balance temporal and spectral resolution, the number of cycles for each wavelet was set as a function of frequency (n=frequency/2). Values were then transformed into a decibel scale relative to a baseline period (−1.9 s to −1.7 s relative to the button press). To assess the power spectra modulation over time and sensors, a one-sample spatio-temporal-spectral cluster-based permutation test (MNE-Python toolbox with default parameters) was applied on the entire time window. As this analysis did not reveal any significant difference, subsequent analyses were conducted on data with lower dimension (spatio-temporal on averaged frequency band, or temporo-spectral on single sensor).

For subsequent analyses the normalized power was averaged within the alpha (8–12 Hz) and beta band (13–30 Hz) and then averaged across trials for each condition (human vs. computer). In each frequency band, group-level statistical significance was assessed by comparing the mean decibel differences between conditions across participants using a one-sample spatio-temporal cluster-based permutation test (MNE-Python toolbox with default parameters) applied on the entire time window.

Finally, to examine the evolution over time of the entire alpha and beta power spectra, analysis was restricted to a single representative sensor over the motor cortex (left-central 52). Within this channel, a one-sample spectro-temporal cluster-based permutation test (MNE-Python toolbox with default parameters) was used to compare the mean decibel differences across the full time-frequency grid (8-30 Hz) between conditions.

Finally, to evaluate the relationship between the magnitude of the RF, alpha ERD, and beta ERD and the decoding performance of Temporal and Riemannian modules, participants were divided into two groups (n = 9 per group) according to decoding performance within the time window of interest. We tested whether participants with a high decoding performance in the Temporal (resp. Riemannian) domain had larger RF (resp. ERD). A subset of sensors were selected for these analyses corresponding to the sensors identified in the permutation tests of the spatio-temporal univariate analysis. Group differences were assessed over time using a two-sample spatio-temporal cluster-based permutation test implemented in the MNE-Python toolbox. Analysis windows were the same used in the univariate analysis section (RF: −2 s to 0 s; ERDs: −2 s to 1 s). For the permutation procedure, the default one-way ANOVA F-statistic was replaced with an independent-samples t-statistic (threshold = 0.05, degrees of freedom = 16). To further assess the relationship between RF and ERD amplitudes and decoding performance, Spearman correlations were computed within the time window of interest. For this analysis, the magnitude motor preparation markers were averaged over the −300 ms to −250 ms interval.

#### Statistics

The significance level was set at α = 0.05 for all the statistical tests. Correction for multiple comparison was performed using the Holm-Bonferroni method and permutation test for clustering approach. To evaluate the statistical sensitivity of our final sample size (n = 18 participants), a sensitivity analysis was performed in GPower (v3.1.9.7). With *α* = 0.05 and power (1 − *β*) set to 0.95, our sample size was sufficient to detect effect sizes as small as Cohen’s *d* = 0.8 (one-tailed paired *t*-test). This confirmed that our study was adequately powered to detect effects smaller than those previously reported in motor preparation decoding paradigms (*d* = 2.3; (Schultze-Kraft et al., 2016)).

## Code Accessibility

All custom analysis scripts used to process the data, generate the figures and statistical results in this manuscript are publicly available on GitHub at https://github.com/RomaricThiboud/Detecting_movement_preparation_with_hybrid_model.git. Raw data required to reproduce the main findings will be made publicly available at the time of the publication.

## Results

### Behaviour

Behaviour in the task was mainly used to evaluate if “computer” trials and “human” trials differ significantly in terms of button press time. The experiment was designed to minimize the difference between distributions. Group-level analysis revealed significant differences in Gamma distributions between “human” trials (mean: scale= 0.441 ± 0.113 ; shape= 5.055 ± 1.720) and “computer” trials (mean: scale= 0.585 ± 0.167 ; shape= 4.044 ± 0.664). Specifically, both scale (one-sample t-test, t=-3.391, p=0.003) and shape parameters (one-sample t-test, t=2.967, p=0.009) differed significantly across conditions, “human” trials having a smaller scale but larger shape compared to “computer” trials (**Fig. 3**). Nonetheless, the overlap between the 2 distributions was high at 93.6%.

**Figure 3:**
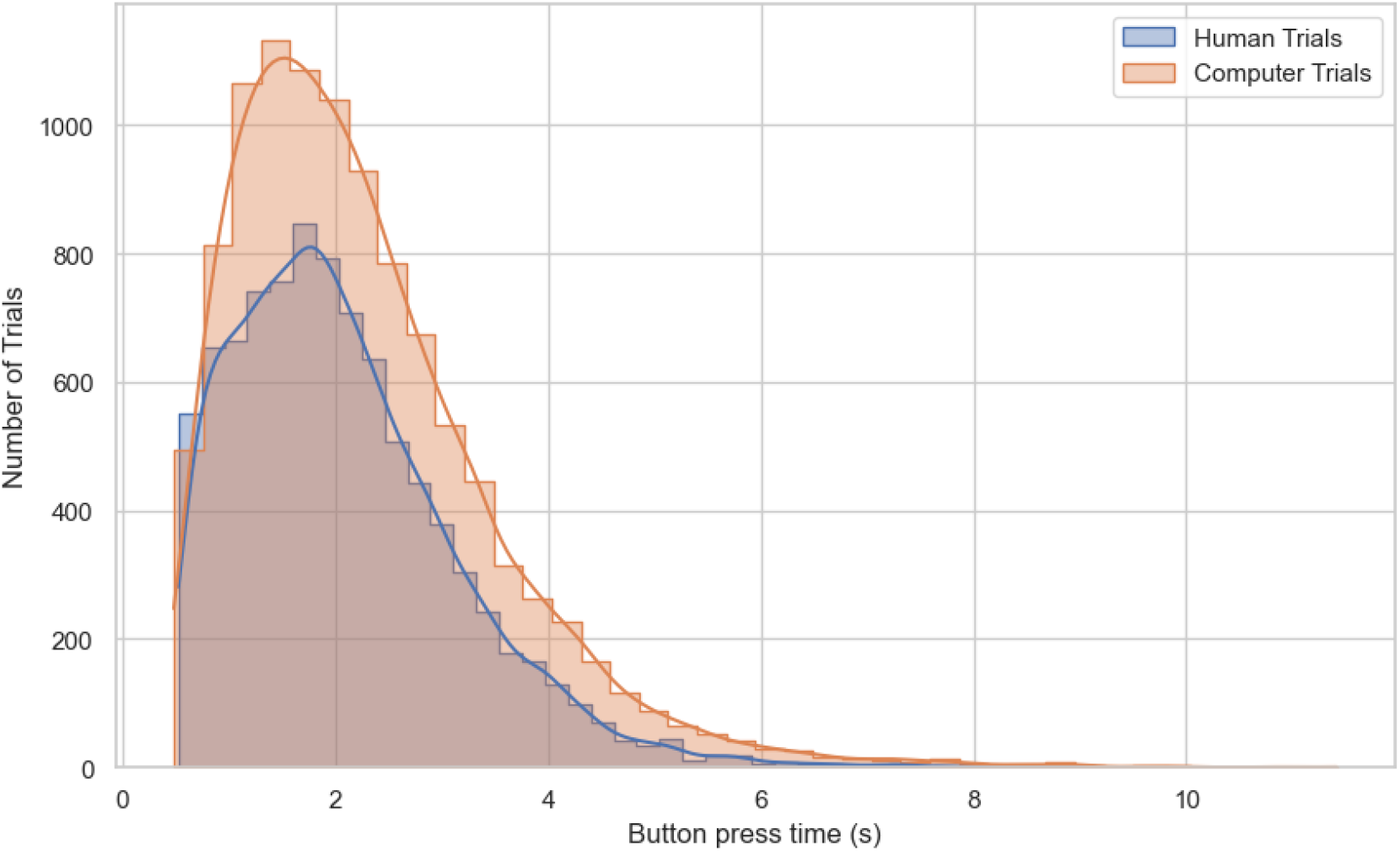
Distribution of button press times for computer trials (orange) and human trials (blue). Data include all 18 participants’ RTs, with human trials shorter than 500 ms excluded (the gamma distribution used for generating computer trials was truncated at 500 ms by design). Solid lines represent gamma function fits to the respective distributions (human: scale=0.44±0.11 ; shape=5.05±1.72 ; computer: scale=0.58±0.17 ; shape=4.04±0.66).

### Multivariate

All the classifiers reached significant above chance decoding (***Fig. 4.A***, p*cluster* < 0.05, one-sample permutation cluster test) at least 850 ms before the button press, the worst performing model in this regard being the gradient boosting tree applied on PCA-extracted features. Across all methods, classification performance (ROC AUC) showed a significant increase over time (all p-values < 0.05, ***Table S1***), reaching a peak at the time of the button press. Note the smaller range of Rho values of the HYDRE classifier compared to the other tested methods. The HYDRE classifier consistently outperformed all the other approaches, with above chance decoding starting at the beginning of the tested window (−2000 ms), a group-averaged ROC AUC of 64% over the time window of interest and an averaged peak ROC AUC of 76% at the time of the button press. The HYDRE classifier showed significantly higher decoding performance (***Fig. 4.B***, pcluster<0.05, one sample permutation cluster test) compared to all the classifiers over a 450 ms long time window (−500 ms to −50 ms before the button press) that encompasses our time window of interest. However, an analysis of the ROC AUC variance at the group level between HYDRE and the other classifiers revealed no significant difference (all p-values > 0.05, ***Table S2***).

**Figure 4.**
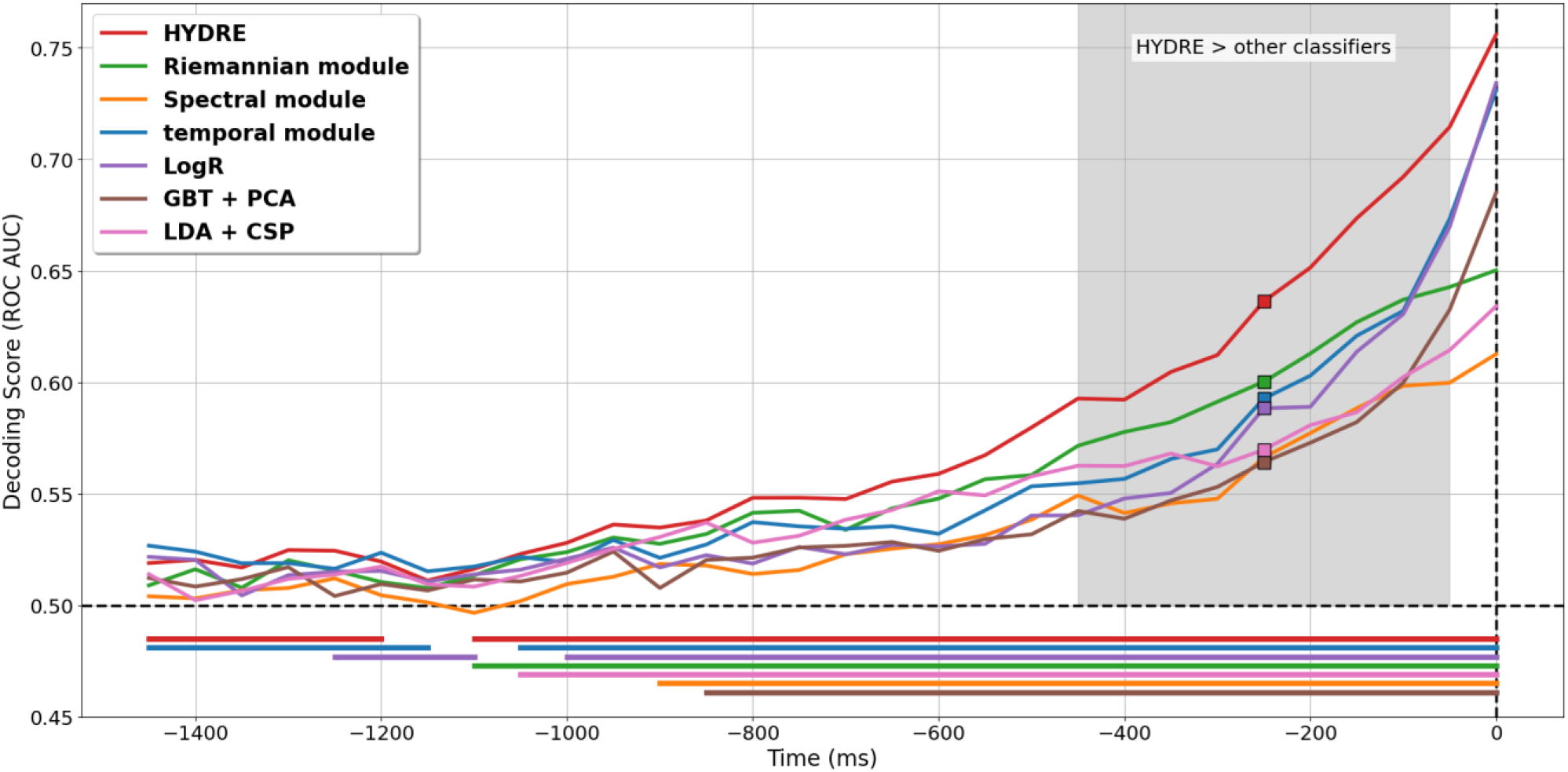
Temporal decoding performance of HYDRE and comparison models. Decoding performance over time for HYDRE and competing classification methods. Each curve represents the group-averaged area under the receiver operating characteristic curve (ROC AUC) across time windows for each classifier. Horizontal colored bars indicate time intervals with above-chance decoding (cluster-based permutation test, p < .05). Square markers indicate decoding performance within the window of interest (300–250 ms before button press). Chance level (AUC = 0.5) is shown as an horizontal black dashed line, vertical dashed line represents the button press time. Grey shaded areas indicate time intervals in which HYDRE significantly outperforms all the other tested classifiers, as assessed using cluster-based permutation tests.

On the time window of interest, the median of ROC AUC scores for the HYDRE, Temporal module, Riemannian module and Spectral module were 0.64, 0.59, 0.60 and 0.57, respectively (***Fig. 5.E***). A Friedman test indicated a statistically significant difference in performance across the three classifiers (χ2 = 23.13, p = 3.79e-5). Subsequent post-hoc analysis using the Nemenyi test revealed that HYDRE outperformed all its constituent modules (***Fig. 5.E***, Temporal p = 0.023, Riemannian p = 0.034, Spectral p = 1.1e-5). No other pairwise comparisons reached statistical significance. The best module was the temporal one for 8 participants, Riemaniann for 9 participants and spectral for 1 participant. On the same time window, the ROC AUC of all the modules were uncorrelated (***Fig. 5.A*, *Fig. 5.C***, p > 0.05) except for the Riemannian and Spectral module (***Fig. 5.B***, *Rho*=0.62, p = 0.018). A strong correlation also exists between the best performing module by participant and the decoding performance of the HYDRE classifier (Rho=0.9, p = 1.6e-6)(***Fig. 5.D***).

**Figure 5.**
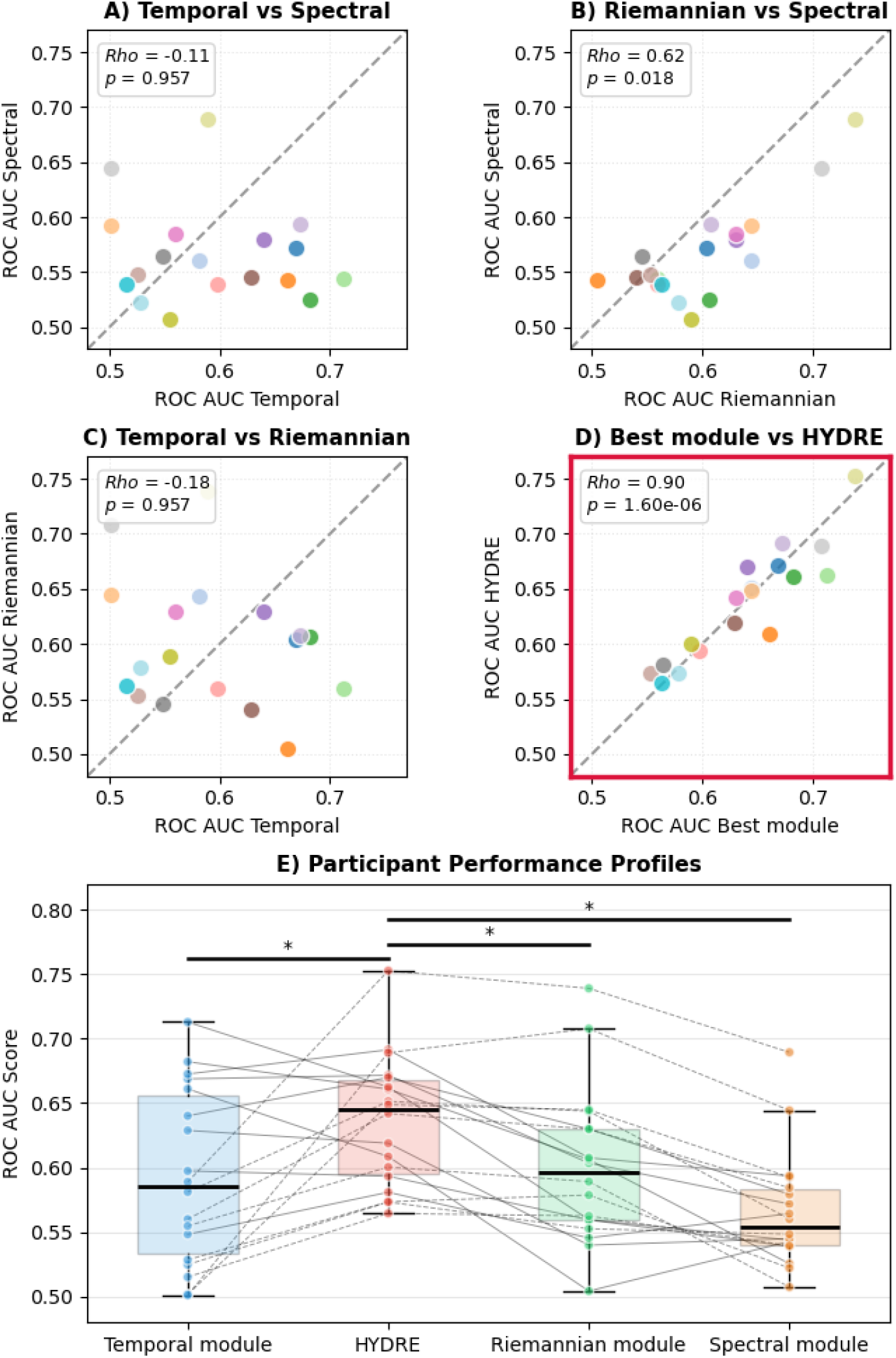
Decoding performance (ROC AUC) comparison between the HYDRE classifier and its constituent modules during the time window of interest. **(A–C)** Pairwise comparison of decoding performance between different modules within the time window of interest (300–250 ms before button press). **(D)** Comparison between the best-performing module for each participant and the HYDRE classifier. In **(A–D)**, the gray dashed line indicates the identity line (y = x), corresponding to equal performance between methods. **(E)** Boxplots showing the distribution of ROC AUC values across participants for the three modules and the HYDRE classifier within the time window of interest. Colored dots indicate performances per participant in the different classifiers. The continuous and dotted lines indicate, respectively, individual data for participants with higher performance for temporal decoding or Riemannian decoding. Black horizontal bars marked with a star indicate significant group-level differences in decoding performance (p < 0.05). During the time window of interest, by aligning its prediction on the best predicting module per participant, the HYDRE model outperforms its composing modules at the group level.

On the time window of interest, the HYDRE classifier presented a better true positive rate on false positive rate ratio compared to the other classifiers on a wide range of decision thresholds (***Fig. 6***). The optimal thresholds, which maximize the combination of sensitivity and specificity and were found using Youden’s J statistic, were liberal for all models, i.e., the “motor preparation” label was selected for probabilities exceeding a threshold lower than 0.5 (mean=0.42±0.024; with a group average optimal threshold of 0.45±0.008 for the HYDRE classifier).

**Figure 6:**
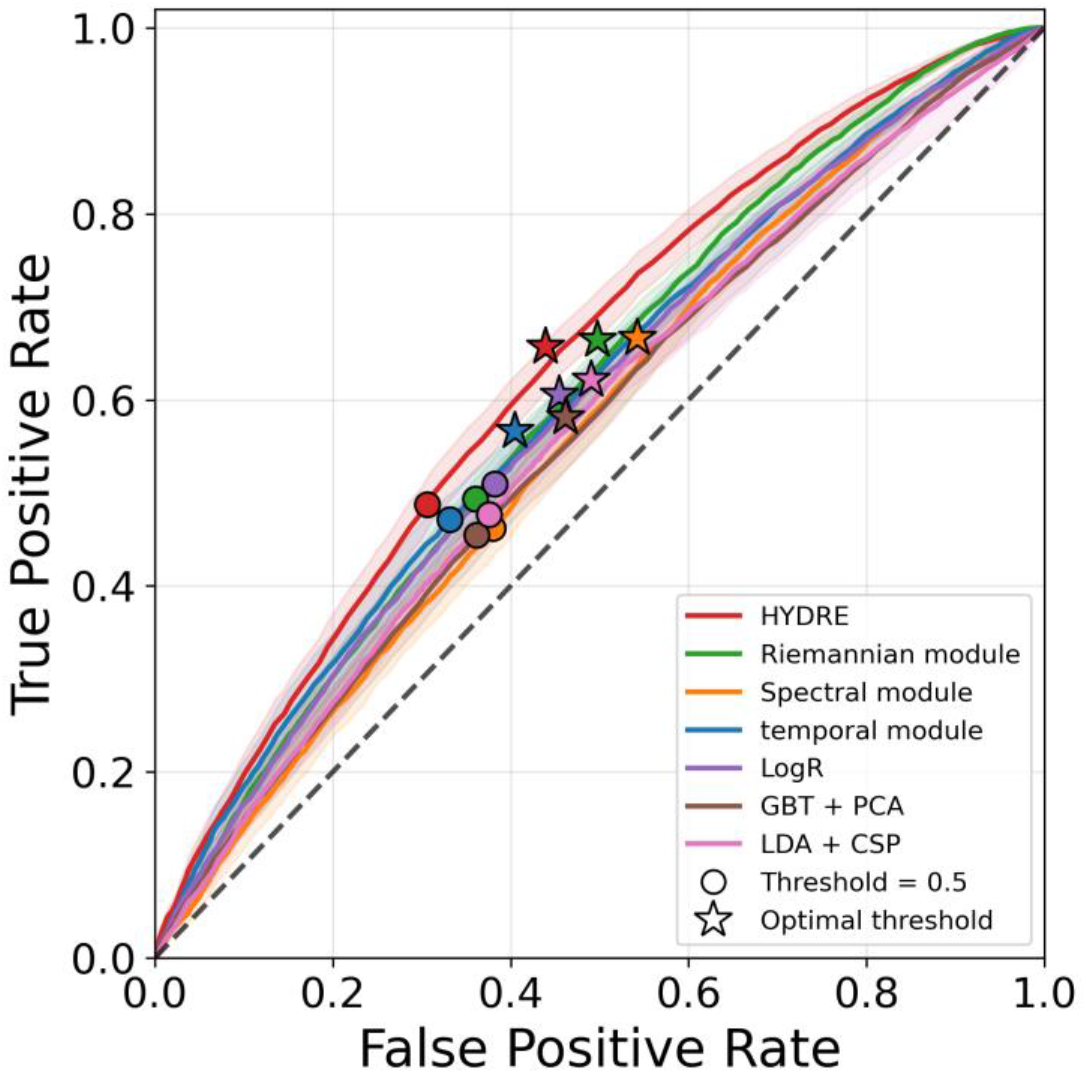
ROC curve of the tested classifiers at the time window of interest (−300 ms to −250 ms before the button press). The grey dotted line represents the chance level. Circles indicate the operating point obtained with a decision threshold of 0.5. Stars indicate the operating point corresponding to the optimal threshold maximizing the Youden index (i.e., both the sensitivity and specificity). On the time window of interest, HYDRE classifier presented a better true positive rate on false positive rate ratio compared to the other classifier for all the possible decision thresholds.

**Figure 7** shows the group-level time course of the Cohen’s kappa coefficient, averaged across the three possible pairs of modules: Temporal-Riemannian, Temporal-Spectral, and Riemannian-Spectral. The group-level average kappa showed a significant increase over time (average Rho = 0.58, p-value = 3.81e-6), starting around an averaged kappa value of 0.01 and peaking at 0.1 coincident with the button press. Throughout the time window of interest, the inter-module agreement remained significantly above zero and below 1 (t-statistic = 6.19 ; p-value = 9.91e-6 ; 95% CI [0.04, 0.08]), meaning that they were neither independent, nor redundant but show some level of complementary.

**Figure 7:**
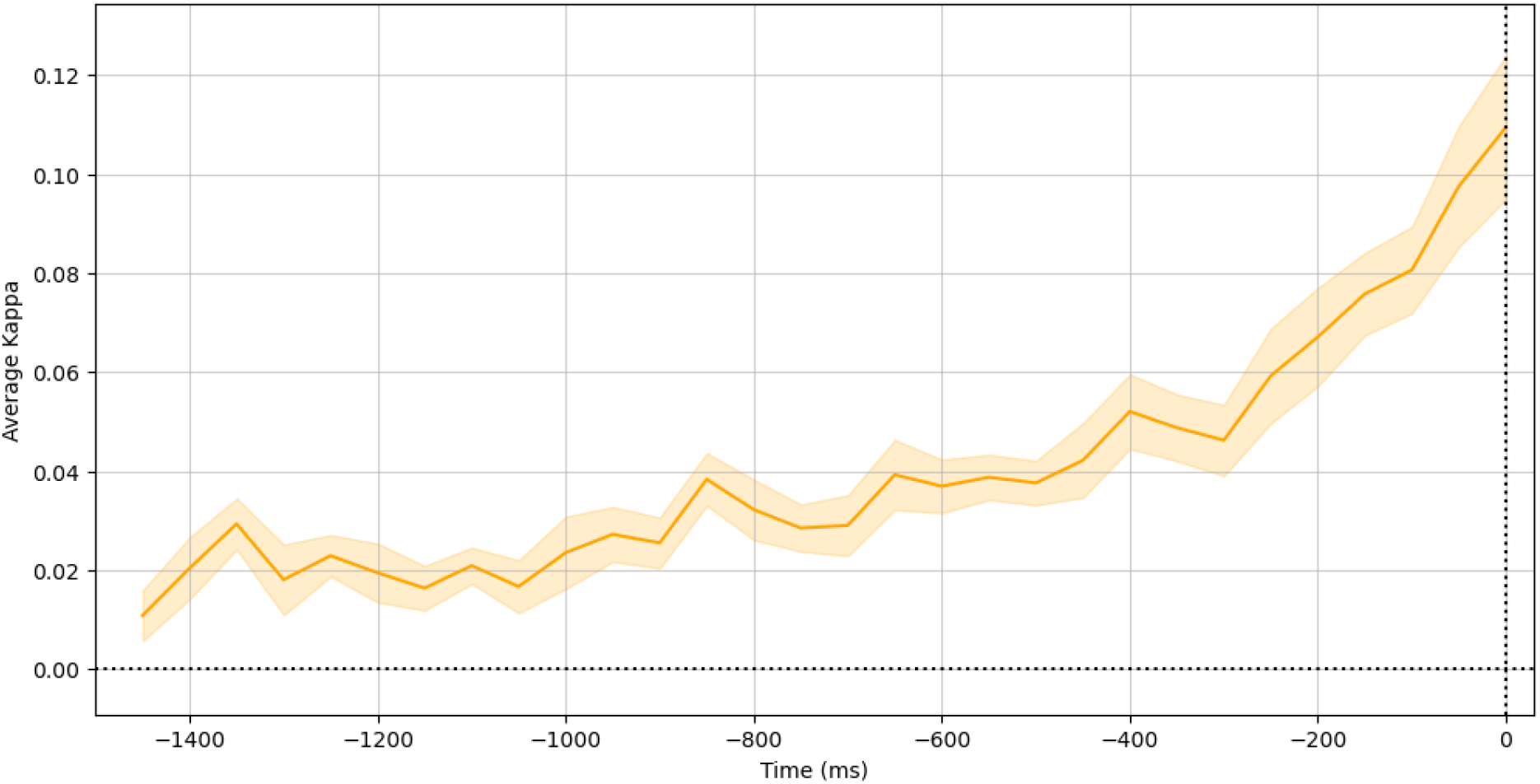
HYDRE modules present a low agreement level (Cohen’s kappa) across time. The light orange shaded area represents the SEM at the group level. HYDRE’s modules are neither independent nor redundant but show some level of complementary.

### Univariate

#### Temporal results

The cluster permutation test performed on the temporal domain over the last 2 seconds leading to the image offset revealed the presence of a RF characterised by a significant difference between “human” and “computer” trials (*pcluster* = 0.001) starting 792 ms before the button press (***Fig. 8.A, bottom***). Over this time window, the cluster spread over the left temporal sensors and bilateral centro-parietal sensors (***Fig. 8.A, left***). The cluster started over centro-temporal sensors on the hemisphere contralateral to the movement, this pattern becoming increasingly bilateral and located over centro-parietal sensors the closer it gets to the button press (***Fig. 8.A, top***).

**Figure 8.**
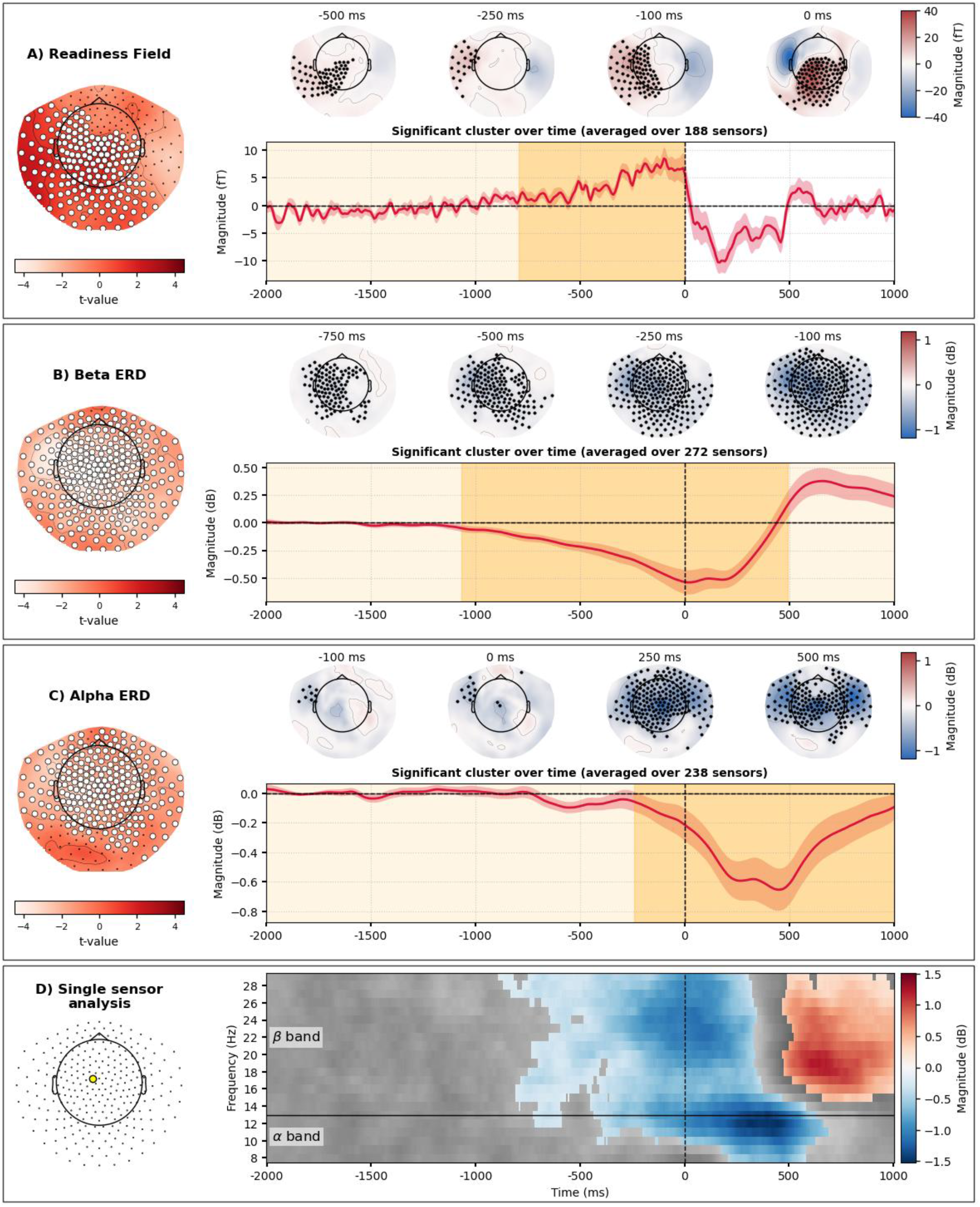
Motor preparation signals in temporal (RF) and spectral (ERD) domains. (**A–C**) Spatio-temporal cluster-based permutation test results. The time series plots show the time course of the signal difference between human and computer trials for the readiness field (RF) (A), the beta ERD (B) or the alpha ERD (C), averaged across all sensors belonging to significant clusters; shaded regions indicate SEM. Light orange shading denotes the full tested time window, and dark orange highlights the significant cluster window (p_cluster_ < 0.05). The topographical maps on top of the time series show topographies of the grand-average difference at several time points during motor preparation and execution; black markers indicate the temporal extent of significant clusters at these specific time points. The topographical map on the left of each panel shows the topographical distribution of t-values; sensors included in the significant cluster at any time point are highlighted in white. (**D**) Time– frequency representation of power differences (dB) between human and computer trials at a representative left-central sensor (MLC-52). The horizontal black line marks the alpha–beta boundary, and the vertical dashed line indicates the button press. Colored regions denote time– frequency points reaching significance (p**_cluster_** < 0.05) based on a one-sample cluster-based permutation test. Trials completed by the participant’s button press elicited significant motor preparation markers in both temporal and spectral domain compared to computer trials.

#### Spectral results

Cluster permutation test over time, frequencies and sensors did not yield any significant result (p*cluste*r>0.05). In the following section we focus on spatio-temporal analysis on averaged frequency band and spectro-temporal single sensors analysis. The cluster-based permutation test performed on the beta-band (13-30 Hz) revealed the presence of a beta-ERD with a significant difference between “human” trials and “computer” trials (*pcluster*= 0.001). This effect was characterized by a decreased beta-band power starting at least 1067 ms before until 492 ms after the button press (***Fig. 8.B, bottom***). Over this time window, the cluster spread over most of the sensors (***Fig. 8.B, left***). The cluster started over central sensors in the hemisphere contralateral to the action (topomap −750ms) before spreading bilaterally (***Fig. 8.B, top***). A second cluster based permutation test performed on the alpha range (8-12 Hz) showed the presence of an alpha-ERD with a significant difference between the 2 types of trials (*pcluster* = 0.005). With a significant decrease of alpha power starting closer to the button press (−242 ms) compared to the cluster in the beta band (−1076 ms) and extending more than 1s after the button press (***Fig. 8.C, bottom***). Over this time window, the cluster spread over most sensors except bilateral occipital sensors (***Fig. 8.C, left***). The cluster started over the left temporal sensors in the hemisphere contralateral to the movement before spreading over bilateral centro-temporal sensors (***Fig. 8.C, top***). Time-frequency analysis of a representative sensor over motor area (MLC-52) revealed dynamics similar to the average frequency band analysis (***Fig. 8.D***). The cluster-based permutation test identified two significant clusters. The first cluster (*pcluster* = 0.001) spanned from −891 ms to 683 ms relative to the button press, encompassing both the alpha- and beta-ERDs. This cluster was characterized by a significant decrease in power in “human” trials compared to “computer” trials. Specifically, alpha-ERD was initially restricted to the high alpha range (>10 Hz) before broadening to include lower alpha frequencies following the button press. Furthermore, the beta-ERD decrease emerged earlier (−891 ms) than the alpha-power decrease (−716 ms). A second cluster (*pcluster* = 0.026) was observed exclusively in the beta band from 466 ms to 1000 ms post-button press, showing an increase in power for “human” versus “computer” trials corresponding to the Event Related Resynchronisation.

Participants for whom the Temporal module showed the lowest performance over the time window of interest exhibited significantly reduced RF magnitude across five brief time clusters spanning −617 ms to −133 ms (***Fig. 9.D*** ; pcluster < 0.05), compared to participants with the highest Temporal module performance. In a similar manner, participants with the poorest Riemannian module performance showed significantly reduced alpha- and beta-ERD magnitudes (***Fig. 9.E–F*** ; pcluster < 0.05) relative to those with the best performance. Group differences in beta-ERD were observed in a cluster extending from −250 ms to 208 ms, while alpha-ERD differences spanned a broader interval from −233 ms to 375 ms around the button press. Within the time window of interest, RF magnitude was positively correlated with Temporal module decoding performance (***Fig. 9.A*** ; Rho = 0.62, p = 0.017), whereas beta-ERD (***Fig. 9.B*** ; Rho = −0.53, p = 0.024) and alpha-ERD (***Fig. 9.C*** ; Rho = −0.55, p = 0.024) magnitude was negatively correlated with Riemannian module decoding performance.

**Figure 9.**
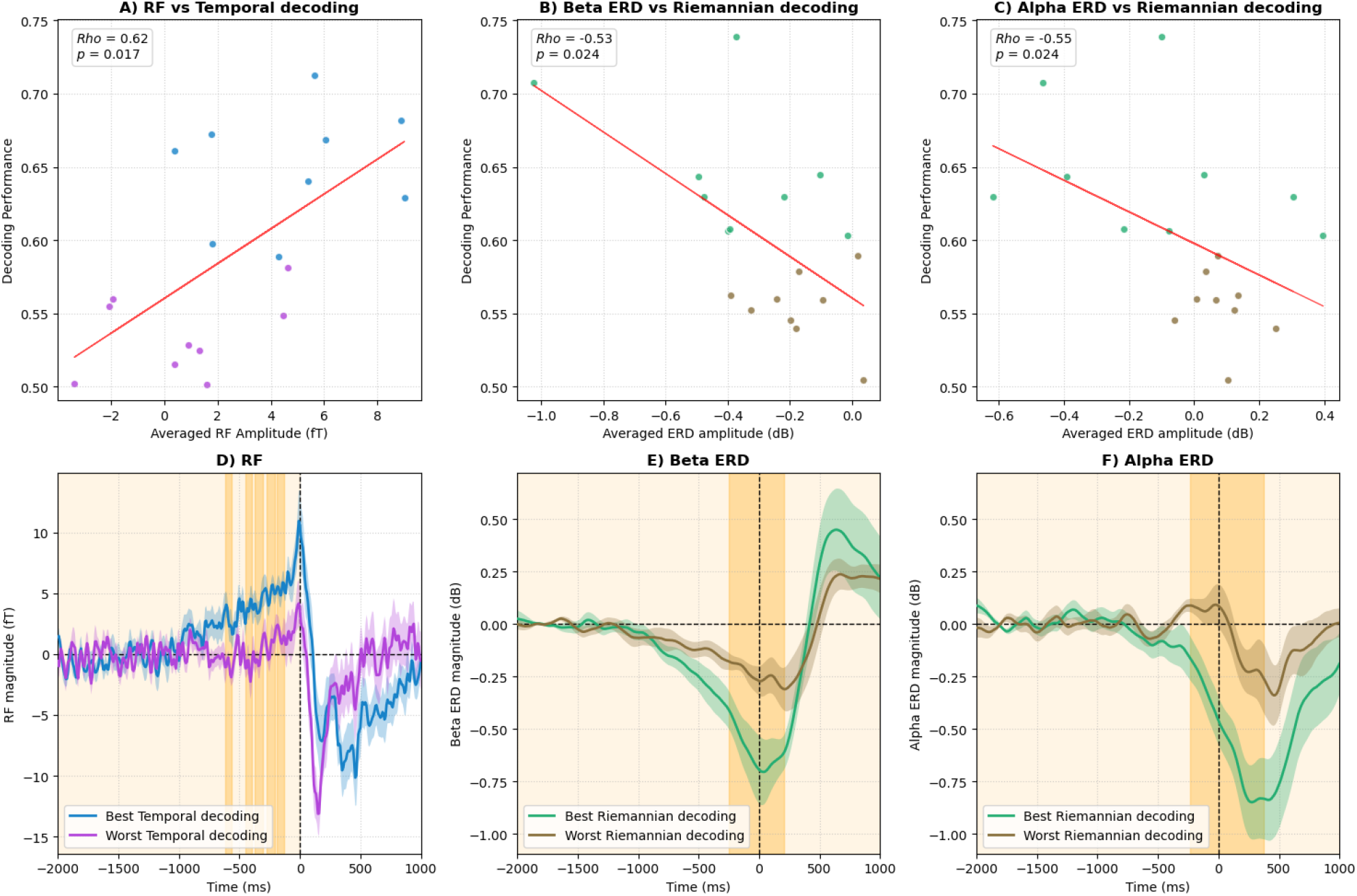
Relationship between the magnitude of motor preparation signal and decoding performance. **(A–C)** Correlations within the time window of interest (300–250 ms before button press) between Readiness Field (RF) amplitude and temporal-module decoding performance (**A**), and between alpha/beta Event-Related Desynchronisation (ERD) and Riemannian-module decoding performance (**B-C**). Significant correlations (Holm-corrected) are indicated by red regression lines. (**D–F**) Time courses of motor preparation marker amplitudes split by participant decoding performance. The upper half (best performers; n = 9) is shown in blue (**D**) or green (**E-F**) and the lower half (worst performers; n = 9) in purple (**D**) or brown (**E-F**). Shaded areas indicate SEM. Light orange shading denotes the full tested interval for the two-sample spatio-temporal cluster-based permutation test, and dark orange shading indicates time periods of significant differences (p**_cluster_** < 0.05). Decoding performances of the temporal and riemannian modules is predictive of the RF and ERD magnitude respectively.

## Discussion

This study aimed at providing more efficient tools to decode motor preparation signals preceding voluntary, self-paced actions. To this aim, we developed a new classification method, HYDRE, that combines features from the temporal, spectral and Riemannian domains in an ensemble classifier. Across the seven classifiers tested, significant above-chance decoding was achieved up to 1.5 seconds before movement, albeit with modest performance. Performance of the classifiers rises relatively close to the button press, the earliest classifier (HYDRE) crossing 60% ROC AUC score 400ms before the button press. We found that the HYDRE classifier attains the highest decoding performances (64% ROC AUC score) on a time window of interest (from −300 ms to −250 ms before the button press). Crucially, an evaluation of the HYDRE classifier revealed weak redundancy of its constituent modules, with a strong correlation between the score of the best performing modules by participant and the decoding performance of HYDRE. In addition, univariate analysis revealed presence of RF and alpha/beta ERD when contrasting “human” and “computer” trials. The magnitude of these motor preparation markers are correlated to the decoding performances of the Temporal module for RF and the Riemannian module for ERD.

Here we showed that the HYDRE classifier was able to decode motor preparation significantly above chance as early as 1500 ms before the button press. This finding aligns with the report by Bai et al. (2011), whose BCI detected motor preparation even earlier than 1.5 seconds before movement initiation, although the majority of their classifier’s predictions occurred in the final 600 ms. Similarly, Schultze-Kraft et al. (2016) have shown silent prediction of their BCI (i.e prediction without triggering consequences) up to 1 second before movement initiation with an increasing number of predictions in the last 300-200 ms before movement initiation.

Evaluation of the HYDRE classifier immediately prior to the button press showed higher performance (68.6% accuracy) than the classifier proposed by Rodrigo et al. (2011). Their model was designed to distinguish between a rest period and motor preparation for arm reaching toward one of eight target locations, achieving 64.0% accuracy. Regarding the trajectory of motor preparation decoding over time, we observed a similar pattern as Ody et al. (2023) with a slow rise of decoding performance before the action followed by a steeper increase. In our study, the HYDRE ROC AUC score started its steeper rise earlier, around 600ms before the button press (***Fig. 4***). This difference can be explained by the combination of features from different domains, the Riemannian module performance over time being more linear and outperforming the temporal module up until 100ms before the button press.

In the window time of interest (300–250 ms pre-button press), HYDRE’s higher ROC AUC compared to competing classifiers, paired with a strong correlation to its best constituent module per participant, indicates that ensembling modules significantly improved decoding performance across participants. However, the complementary analysis indicates that the gain realized by the ensemble did not exceed the performance of the best performing module for each participant. Indeed the Cohen’s Kappa value slightly above 0 indicates very weak redundancy but no real complementarity (***Fig. 7***). Ieracitano et al. (2021) reported an averaged ensemble gain (i.e. accuracy gain between the accuracy of the best domain and the accuracy of the ensemble) at the group level of 7.37% ± 3.72% accuracy. But some differences can be noted, (1) the combination of continuous wavelet transforms and convolutions could better capture evolving structures and transient events across time, (2) decoding was performed in source space that could have improved discriminability between rest and movement preparation. Another study, methodologically closer to ours, demonstrated a gain of accuracy by combining functional connectivity and covariance-based features within an ensemble of linear classifiers that improves right/left hand motor imagery classification (Corsi et al., 2022). While the authors concluded that the feature diversity mitigated inter/intra individual variability, thus improving performance, our findings offer a more nuanced view. While the HYDRE classifier improved overall, decoding performance at the group level, it did not significantly reduce performance variance across participants. This suggests that while multi-domain information provides complementary benefits, it is not sufficient alone to solve the broader challenge of “BCI illiteracy” (Ahn & Jun, 2015).

Interestingly, the performance of modules trained on Riemannian and spectral features was strongly correlated. This likely reflects that both feature families derive from the same second-order statistics of the MEG signal. Riemannian tangent-space features are computed from covariance matrices, whose diagonal elements represent channel variances and whose off-diagonal elements capture cross-channel covariation (Congedo et al., 2017), while band power corresponds to signal variance within specific frequency bands. Oscillatory activity that drives band power differences therefore also shapes the covariance structure used by Riemannian methods, making their predictions correlated. Conversely, the visible but non-significant performance difference between the two modules may reflect differing robustness to preprocessing: although artifact rejection is generally beneficial, Riemannian models remain robust without it, unlike models based on spectral power (Sabbagh et al., 2020).

We also observed that the best module varies across participants. Furthermore, no module outperformed all the other ones in average decoding performance. (***Fig. 5.E***). These results suggest that depending on the individual, the availability of relevant brain activity for motor preparation is more prominent in one or the other domain. Indeed, despite consistent group-level motor preparation markers, univariate analysis revealed substantial inter-individual variability that may originate from several factors. First, during the task, participants were instructed to press the button whenever they wanted, but did not receive any instructions regarding the speed or force of their movement. As movement parameters have been shown to affect the magnitude and/or latency of the RF (Shibasaki & Hallett, 2006), variability in these parameters may have contributed to variability in the observed RF. On the more cognitive side, the attentional demands of the task of the present study was low, relying on the repetition of a simple motor action with no requirement for action selection. Under such conditions, participants may progressively shift toward a more automatic mode of execution, reducing the engagement of higher-order motor planning processes. Consistent with this interpretation, recent work has suggested that the amount of neural resources allocated to motor planning directly influences the magnitude of ERD (Pierrieau et al., 2023). Variability in ERD magnitude may therefore reflect differences in the degree of motor planning engaged by individual participants. Finally, variability could come from personality traits of participants; notably, it has been shown that impulsivity is positively correlated to the magnitude of ERD (Tzagarakis et al., 2019) and RF (Rossi et al., 2018). Taken together, differences in movement parameters, motivational state, and intentional engagement may represent important sources of inter-individual variability in the neural signatures of motor preparation observed in the present study.

Our results indicate that averaged motor preparation markers are functionally relevant at the single-trial level, correlating with temporal and Riemannian decoding performance (Fig. 9A-C). Specifically, RF magnitudes between easy- and hard-to-decode participants diverged as early as 600 ms before movement onset. This early relationship supports the traditional view of the RP as an index of motor preparation, while partially contradicting the stochastic accumulator model (Schurger et al., 2012), which cannot easily account for such early divergence via spontaneous fluctuations.

Note that we tested the HYDRE classifier in a challenging task that did not reinforce motor preparation marker as it is a simple (Erdler et al., 2000; Patil & Kochhar, 2025), non rewarding (Schultze-Kraft et al., 2016), self-paced (Salvaris & Haggard, 2014), uncued (Ody et al., 2025) task. Nonetheless HYDRE classifier presented the best performance among comparative classifiers, but it remains to be tested if it is also the case in paradigms that elicit clearer motor preparation markers. Moreover, due to the continuous nature of the task the experiment lacked a distinct intertrial interval with events frequently overlapping. Furthermore, in “computer” trials, participants may have already begun preparing a movement before successfully aborting it. As a result, neither “human” nor “computer” trials reflect purely isolated states of motor preparation or rest. On the one hand, this could have decreased decoding performances of HYDRE. On the other hand, it is closer to real-life applications, where the BCI system will receive contradictory or noisy signals. Lastly, the HYDRE model was evaluated exclusively offline. Translating this approach to real-time deployment requires selecting an optimal operating threshold (***Fig. 6***), ensuring the model generalizes well across the motor preparation window, and keeping prediction latency low.

In conclusion, integrating temporal, spectral, and Riemannian features within an ensemble classifier (such as HYDRE) significantly improves the decoding of motor preparation. By leveraging the complementarity of these domains, this approach offers a robust solution for motor BCIs to overcome the high inter-subject variability typically observed in motor preparation markers.

## Acknowledgments

This work was supported by the Impulsion Grant from IDEXLYON (Initiative d’Excellence de l’Université de Lyon) and ANR MyAct (ANR-23-CE28-0009).

We would like to thank Salim KLIBI and Sébastien DALIGAULT for data acquisition, Sonia ALOUCHE and Cecile TROUBA for administrative support.

Generative AI language tools were utilized during manuscript preparation solely for language editing, grammar improvement, and text rephrasing. All generated suggestions were critically reviewed, edited, and finalized by the authors, who retain full responsibility for the final text.

---

**Table S1:** Decoding performances increase over time for all tested classifiers. Note the smaller range of Rho values for the HYDRE classifier.

| Classifier | Averaged Rho | Std. Rho | Range Rho | W-statistic | Corrected p-value (Holm-Bonferroni) |
| --- | --- | --- | --- | --- | --- |
| **HYDRE** | 0.86 | 0.08 | [0.67 - 0.96] | W > 171.0 | p<10<sup>-4</sup> |
| **Riemannian module** | 0.78 | 0.17 | [0.35 - 0.96] | W > 171.0 | p<10<sup>-4</sup> |
| **Spectral module** | 0.69 | 0.19 | [0.22 - 0.95] | W > 171.0 | p<10<sup>-4</sup> |
| **Temporal module** | 0.64 | 0.24 | [0.09 - 0.96] | W > 171.0 | p<10<sup>-4</sup> |
| **Logistic Regression** | 0.61 | 0.25 | [0.11 - 0.96] | W > 171.0 | p<10<sup>-4</sup> |
| **GBT + PCA** | 0.60 | 0.28 | [0.06 - 0.95] | W > 171.0 | p<10<sup>-4</sup> |
| **LDA + CSP** | 0.71 | 0.17 | [0.31 - 0.95] | W > 171.0 | p<10<sup>-4</sup> |

**Table S2:** No significant difference in decoding performance variance at the group level on the time window of interest.

| Comparison | Variance<br>HYDRE | Variance<br>Other | t-statistic | Corrected p-value<br>(Holm-Bonferroni) |
| --- | --- | --- | --- | --- |
| **HYDRE / Riemannian module** | 0.00257 | 0.0035 | -0.0095 | 1.00 |
| **HYDRE / Spectral module** | 0.00257 | 0.0020 | 0.0006 | 1.00 |
| **HYDRE / Temporal module** | 0.00257 | 0.0046 | -0.0021 | 1.00 |
| **HYDRE / Logistic Regression** | 0.00257 | 0.0029 | -0.0004 | 1.00 |
| **HYDRE / GBT+PCA** | 0.00257 | 0.0023 | 0.0003 | 1.00 |
| **HYDRE / LDA+CSP** | 0.00257 | 0.0029 | 0.0003 | 1.00 |

